# mTADA: a framework for analyzing de novo mutations in multiple traits

**DOI:** 10.1101/406868

**Authors:** Hoang T. Nguyen, Amanda Dobbyn, Joseph Buxbaum, Dalila Pinto, Shaun M Purcell, Patrick F Sullivan, Xin He, Eli A. Stahl

## Abstract

Joint analysis of multiple traits can result in the identification of associations not found through the analysis of each trait in isolation. In addition, approaches that consider multiple traits can aid in the characterization of shared genetic etiology among those traits. In recent years, parent-offspring trio studies have reported an enrichment of *de novo* mutations (DNMs) in neuropsychiatric disorders. The analysis of DNM data in the context of neuropsychiatric disorders has implicated multiple putatively causal genes, and a number of reported genes are shared across disorders. However, a joint analysis method designed to integrate de novo mutation data from multiple studies has yet to be implemented. We here introduce multi pi e-trait TAD A (mTADA) which jointly analyzes two traits using DNMs from non-overlapping family samples. mTADA uses two single-trait analysis data sets to estimate the proportion of overlapping risk genes, and reports genes shared between and specific to the relevant disorders. We applied mTADA to >13,000 trios for six disorders: schizophrenia (SCZ), autism spectrum disorder (ASD), developmental disorders (DD), intellectual disability (ID), epilepsy (EPI), and congenital heart disease (CHD). We report the proportion of overlapping risk genes and the specific risk genes shared for each pair of disorders. A total of 153 genes were found to be shared in at least one pair of disorders. The largest percentages of shared risk genes were observed for pairs of DD, ID, ASD, and CHD (>20%) whereas SCZ, CHD, and EPI did not show strong overlaps In risk gene set between them. Furthermore, mTADA identified additional SCZ, EPI and CHD risk genes through integration with DD *de novo* mutation data. For CHD, using DD information, 31 risk genes with posterior probabilities > 0.8 were identified, and 20 of these 31 genes were not in the list of known CHD genes. We find evidence that most significant CHD risk genes are strongly expressed in prenatal stages of the human genes. Finally, we validated our findings for CHD and EPI in independent cohorts comprising 1241 CHD trios, 226 CHD singletons and 197 EPI trios. Multiple novel risk genes identified by mTADA also had de novo mutations in these independent data sets. The joint analysis method introduced here, mTADA, is able to identify risk genes shared by two traits as well as additional risk genes not found through single-trait analysis only. A number of risk genes reported by mTADA are identified only through joint analysis, specifically when ASD, DD, or ID are one of the two traits examined. This suggests that novel genes for the trait or a new trait might converge to a core gene list of the three traits.

## 1. Introduction

The analysis of multiple traits can help characterize the genetic architectures of complex disorders (Solovieff, et al., 2013). One approach is to meta-analyze results derived from separate single-trait studies (Zhernakova, et al., 2011). However, joint analysis with multiple traits can better accommodate heterogeneity of genetic effects of the same variants or genes across traits (Allison, et al., 1998; Galesloot, et al., 2014). Numerous studies have jointly analyzed two or more traits and successfully identified shared common-variant associations (Giambartolomei, et al., 2014; Pickrell, et al., 2016; Lutz, et al., 2017; Turley, et al., 2018); however, none of these studies has examined rare variation from case-control (CC) data, or de novo variants for which mutations rates should be taken into account. For these rare variants, gene based tests have successfully identified genes associated with different disorders (He, et al., 2013; De Rubeis, et al., 2014; Iossifov, et al., 2014; Nguyen, et al., 2017). Some recent studies have also shown that there are multiple risk genes that are shared between neurodevelopmental disorders (Hoischen, et al., 2014; Li, et al., 2016; Nguyen, et al., 2017), and also with congenital heart disease (CHD) (Homsy, et al., 2015; Willsey, et al., 2018). These results are based on the intersection among the top prioritized genes from each disorder; therefore, reported numbers of genes shared by multiple disorders remain low (Nguyen, et al., 2017; Willsey, et al., 2018). Development of multi-trait rare-variant methods for neuropsychiatric disorders (NPDs) and related disorders will facilitate the understanding of this important aspect of genetic architecture for these phenotypes.

Currently, there is still a limitation in the risk gene identification for a single trait of NPDs and relevant disorders. One reason is that risk gene discovery is underpowered when sample sizes are limited, as well as when relative risks (RRs) are not large (He, et al., 2013; Nguyen, et al., 2017). Multiple risk genes have been reported for developmental disorders (DD), intellectual disability (ID) and autism spectrum disorder (ASD) (De Rubeis, et al., 2014; Lelieveld, et al., 2016; Deciphering Developmental Disorders Study, 2017) thanks to large sample sizes and/or RRs (Nguyen, et al., 2017). However, there are a few risk genes identified for schizophrenia (SCZ) and epilepsy (EPI) because of small RRs and small sample size respectively (Nguyen, et al., 2017). Simply increasing sample size is an expensive solution and might not be feasible for some rare disorders. If the genetics overlap, methods that can leverage the information from one trait to increase power for risk-gene identification for another trait with smaller sample size or RRs could help in obtaining additional genes for these disorders.

Here, we have developed a new statistical model that combines *de novo* mutation information to identify shared and specific risk genes for two disorders. To illustrate the advantage of the new pipeline over its previous single-trait version, we have applied the pipeline to a large data set of different NPDs and CHD (∼ 13,000 parent-offspring trios) and identified shared genes between each pair of these disorders. We have also used this pipeline to identify additional risk genes for each disorder by borrowing the information of other traits. The identification of shared genes is important for understanding the overlapping genetic information of these disorders.

## 2. Methods

### 2.1 The mTADA pipeline

#### 2.1.1 Statistical models in mTADA

We developed the multi-trait Transmission And Denovo Association (mTADA) pipeline to analyze DNMs for any two given disorders using the computational framework of extTADA (Nguyen, et al., 2017) (Table 1). The mTADA pipeline is gene-based and requires input data of the number of *de novo* mutations and mutation rate per gene. If the de novo mutations are stratified on the basis of predicted effect (e.g. ‘missense’, ‘nonsense’, etc.), then each gene-annotation category should have its own mutation rate that reflects the predicted effects of the mutations within. In summary, for each gene, we consider four models *M*_*j*_ (*j* = 0‥3) reflecting four alternative hypotheses: the gene is associated with neither trait (Ho), the first trait only (H_1_), the second trait only (H_2_), or both traits (H_3_). We assume prior probabilities *π*_*j*_*·*, (*j* = 0‥3) for the four models. To build models for these hypotheses, we used single-trait models from TADA (He, et al., 2013); therefore, we first introduce TADA and then mTADA. Like TADA, mTADA divides mutations into different annotation categories (e.g., loss-of-function or missense variants), builds models for these categories, and then integrates information across models to infer results.

**Table 1:**
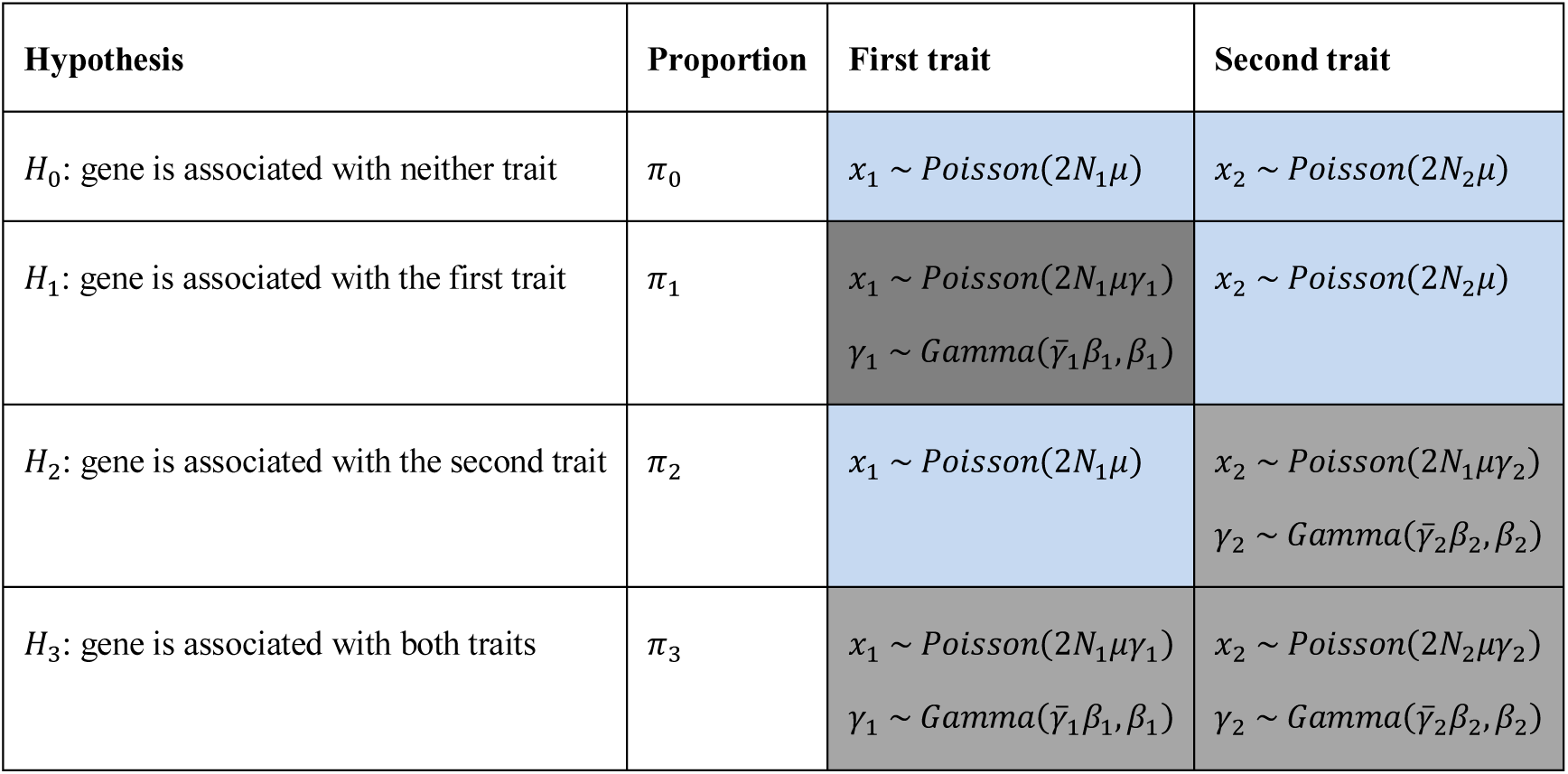
*Statistical models for four hypotheses in mTADA for one category of variants in each trait. For each gene, mTADA assumes that the gene can be in one of four models M*_*0*_*‥M*_3_*. π* _*j*_ (*j* = 0‥3) *is the prior probability of the j*^*th*^ *model. x*_*k*_ *and N*_*k*_ (*k =* 1,2) *are the data and the sample size of the k*^*th*^ *trait. μ is the mutation rate of the gene; γ*_*k*_ *and* 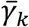 *are the relative risk and mean relative risks of the variant category. C and cells describe models for risk and non-risk genes respectively.*

In TADA, for the single trait *k*, for each variant/mutation category in each gene, all variants are collapsed and considered as one count (*x*_*k*_) with *x*_*k*_*∼Poisson*(*2N*_*k*_*γ*_*k*_), in which *N*_*k*_*,μ* and *γ*_*k*_ are the sample size (family number), mutation rate and relative risk (RR) of the category. For a single trait, TADA compares two hypotheses for each gene: non-risk gene *γ*_*k*_ *=* 1, and risk gene *γ*_*k*_ *>* 1 in which *γ*_*k*_ follows a Gamma distribution: 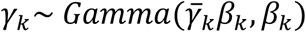 with 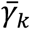 being the mean relative risk (meanRR) across risk genes, and *β*_*k*_ being a dispersion parameter. Thus, TADA implies that a proportion of all genes 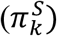 are risk genes. In mTADA, with two traits and four models, we define *γ*_*k*_ as the non-null RR for trait *k* (*k =* 1,2), which we can describe in terms of *γ*_*k,j*_ for trait *k* under the *j*^*th*^ model as *γ*_*1*_ *= γ*_1,1_ = *γ*_1,3_ *>* 1 for the first trait, and *γ*_2_ = *γ* _2,2_ = *γ*_2,3_ > 1 for the second trait (Table 1). Under the non-risk gene models for each trait, *γ* _1,0_ = *γ* _2,0_ *= γ* _2,1_ *= γ* _1,2_ *=* 1.

The likelihood of the data across all *N* genes can be computed as 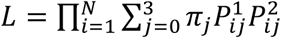 with 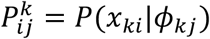, where *x*_*ki*_ and *Φ* _*ki*_ are the *i*^*th*^ gene data and *j*^*th*^ model parameters for trait *k* (*k* = 1,2). In addition, if the data include multiple categories of variants then 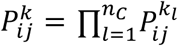 with *n*_*C*_ being the number of categories.

#### 2.1.2 Estimation of the parameters

We used our single-trait pipeline, extTADA, to estimate the proportions of risk genes (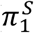 and 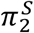), mean relative risks (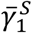 and 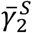) and dispersion parameters (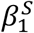 and 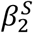) for each single trait (described as the superscript). We used these values inside mTADA: 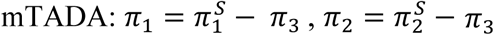, and 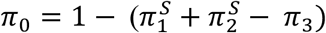 because of 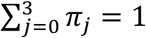. We also assumed that 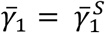 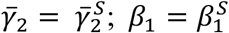 and 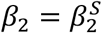. Therefore, we only estimated *π*_3_ inside mTADA. Bayesian models were built using the *rstan* package (Carpenter, et al., 2016). We used Markov Chain Monte Carlo (MCMC) within *rstan* to estimate *π*_3_. We also implemented another option for users to choose the automatic differentiation vibrational inference (ADVI) (Kucukelbir, et al., 2015) to estimate *π*_3_. Convergence was diagnosed by the estimated potential scale reduction statistic (*R*) and visualizing traces of results. The *Locfit* package (Loader, 2007) was used to obtain the mode, credible interval (CI) of *π*_3_. We used the mode as the estimated value of *π*_3_.

#### 2.1.3 Inference of risk genes

For gene *i*, the statistical support for the *j*^*th*^ model is captured by its posterior probability 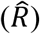, abbreviated as PP0, PP1, PP2 or PP3 for a gene). Inference of risk genes shared by the two traits can be made based on PP3. To summarize the evidence for association with a given trait, we used the sum of posterior probabilities of models including the risk gene hypothesis for that trait (Barber, et al., 2010), i.e. *PP*_*it*_ *+ PP*_*i*3_ for trait one and *PP*_*i*2_ + *PP*_*l*3_ for trait two. In this way, we can clearly see how trait two’s data’s support for risk genes may contribute to support for trait one.

### 2.2 Simulation analysis

#### 2.2.1 Generation of simulated data

We simulated data under the mTADA model in Table 1. A gene was assigned to one of the four groups (four models) by using the probability (*π*_0_*, π*_1_*, π*_2_*, π*_3_). We used 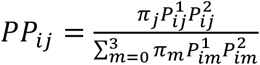 and 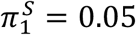 which are approximately equal to ASD, ID and DD results in our single-trait study (Nguyen, et al., 2017). *π*_3_ was simulated with different values between 0 to min 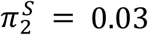, and *π*_0_*, π*_1_, and *π*_2_ were calculated as in the previous section. A range of meanRRs were simulated for each of the two traits. Two mutation categories were simulated for each trait; therefore, there were four meanRRs for the two traits. We simulated 100 values of each combination of *π*_3_, 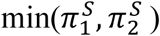 and 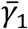 and calculated the mean of these 100 simulation results.

#### 2.2.2 Validation of parameter inference and risk gene discovery

To calculate Type I error, we simulated different combinations of genetic parameters with *π*_3_ = 0. For each PP threshold, if there were at least one significant overlapping gene (PP3 > the PP threshold) then the error was calculated.

We used simulated data to assess the correlation between true and observed *π*_3_ values and between PPs and observed false discovery rates (oFDRs). An oFDR at a PP threshold was defined as the number of false positive genes divided by the number of identified genes. To use mTADA for single traits, for the *i*^*th*^ gene, we calculated *PP*_*i*l_ *+ PP*_*i*3_ and *PP*_*i*2_ *+ PP*_*i*3_ for the first and second trait respectively.

#### 2.2.3 Comparison between single-trait and two-trait pipelines

We used AUCs (area under the Receiver Operating Characteristic (ROC) curve) to compare risk gene classification performance between mTADA and extTADA on single traits. To obtain AUCs, we calculated true and false positive rates for extTADA and mTADA across PP thresholds, and calculated the areas under these ROC curves. We set a threshold of PP=0.8 to compare gene counts between mTADA and extTADA.

### 2.3 Neuropsychiatric disease and congenital heart disease de novo mutation data

We used the DNM data collected by Nguyen, et al. (2017) and CHD data from Homsy, et al. (2015). These data included 356 EPI trios, 5122 ASD trios, 4293 DD trios, 1012 ID trios, 1017 SCZ trios, and 1213 CHD trios. DNMs were annotated and classified into multiple categories as in our previous work (Nguyen, et al., 2017). For NDDs (EPI, ASD, DD and ID) and CFID, we used two categories (Nguyen, et al., 2017): loss-of-function (LoF) and missense damaging (MiD) DNMs. The LoF category included nonsense, essential splice site, and frameshift DNMs defined by Plink/Seq (Fromer, et al., 2014) while the MiD category included DNMs annotated as missense by Plink/Seq and predicted damaging by each of seven methods (Genovese, et al., 2016): SIFT, Polyphen2_FIDIV, Polyphen2_HVAR, LRT, PROVEAN, MutationTaster, and MutationAssessor. For SCZ, we used LoF, MiD and synonymous mutations within DNase I hypersensitive sites (DFISs) because this category showed significant DNM enrichment in SCZ probands (Takata, et al., 2016) and non-null meanRR in extTADA (Nguyen, et al., 2017). Mutation rates were calculated as described by Fromer, et al. (2014) and Nguyen, et al. (2017). extTADA was used to obtain the proportions of risk genes and the meanRR of each category for each disorder. These values were then used as input for mTADA to estimate *π*_3_ and then calculate *PP*_*ij*_ (i = 1‥ N, j = 0‥3) for each pair of traits.

The MCMC algorithm, No-U-Turn Sampler (NUTS), in the *rstan* package was used to estimate *π*_3_. Two independent chains and 5000 steps for each chain were used in the sampling process. Only 1000 samples from each chain were chosen for further analyses.

We used results from new de novo studies to validate mTADA results. New CFH) de novo data include 2,871 probands which consist of 2445 trios (1204 trios are inside the data set of extTADA and used in the primary analysis of this study) and 226 singletons (Jin, et al., 2017). In addition, we used the whole-genome-sequencing (WGS) trio data for EPI (Hamdan, et al., 2017) to validate results of EPI, which include 197 trios not included in our mTADA analyses.

For further in silico validation and characterization genes identified by mTADA, GeNets (Li, et al., 2017) was used to test protein-protein interactions (PPIs). The STRING database (Szklarczyk, et al., 2017) was also used as an alternative source for PPIs. For GeNets, we used results from input genes and direct-connection candidate genes automatically inferred by GeNets. For STRING, we used input genes only to obtain final results. To examine expression information of identified genes, spatiotemporal transcriptomic data were obtained from BRAINSPAN (Miller, et al., 2014), divided into eight developmental time points (four prenatal and four postnatal) (Lin, et al., 2015), and analyzed by hierarchical clustering for developmental trajectories.

To test the significance of the overlap of two gene sets, a permutation approach was used. We chose two random gene sets whose lengths are the same as the two tested gene sets from the background genes (19358 genes from mTADA). This was carried out *N* times (*N* = 10,000 in this study) and the numbers of overlapping genes were recorded in a vector ***m*** P value was calculated as (*length*(***m*[*m*** *> m*_0_]) + 1)/(*length*(**m**) + 1)) in which *m*_0_ is the observed number of overlapping genes between the two tested gene sets.

## 3. Results

Because mTADA is a novel tool that analyzes multiple traits using *de novo* mutation and mutation rate data, we validated mTADA using data simulated under its model (Table 1) and compared gene-identification results with our a single-trait pipeline extTADA.

### 3.1 Results of mTADA on simulated data

We used 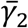 and 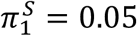 as described in the Method section and simulated different data sets from the combination of different values of *π*_3_ and MeanRRs.

#### 3.1.1 Type I error of shared risk gene identification

We first estimated Type I error for identifying shared risk genes (i.e. associated with both traits). We simulated data with *π*_3_ *=* 0 and tested for shared risk genes using different thresholds of the posterior probability of Model III (PP3). Smaller PP3 thresholds correspond to increased Type I error levels. This error was smaller than 0.05 when PP3 > 0.8 (Figure 1A). Overall, the error decreased when meanRRs or sample sizes increased.

**Figure 1:**
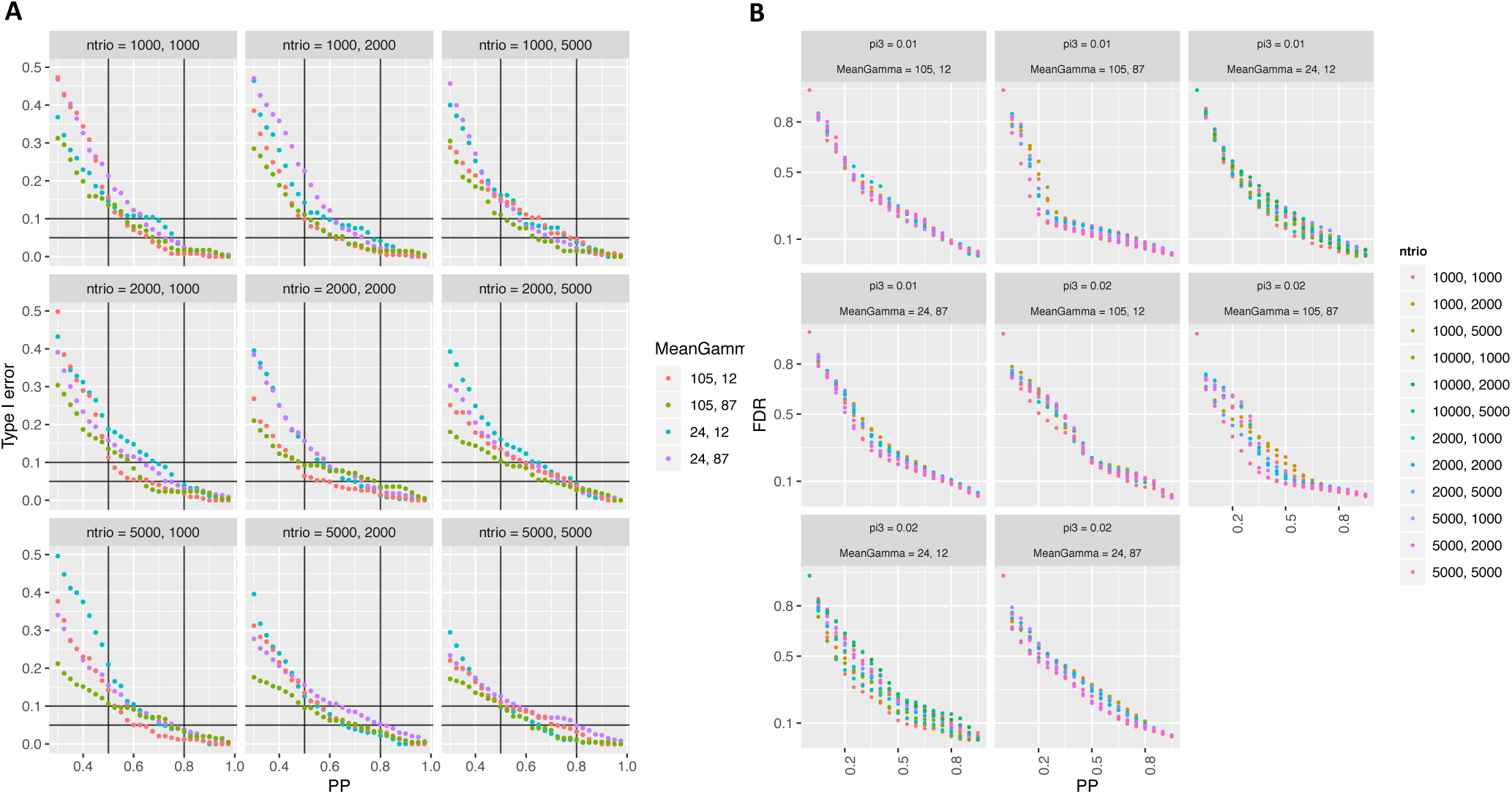
*Validation of shared risk gene identification using mTADA on simulated data. Left panels show the Type I error of the identification of risk genes for both traits: X-axes are posterior probabilities* (*PPs*) *of Model III while Y-axes are Type I errors. Right panels show the correlation between PPs* (*x axis*) *and observed false discovery rates* (*FDRs, y axis*)*. These are for the combination of different sample sizes* (*ntrio*) *and mean relative risks* (*MeanGamma*). ntrio and MeanGamma describe the information of PA>O traits.

#### 3.1.2 Correlations between posterior probabilities and observed false discovery rates

We also calculated the correlation between PPs and observed FDRs (oFDRs) for all situations. Regarding PPS and oFDRs, PP=0.8 and 0.5 were approximately with oFDR=0.1 and 0.25 respectively. Small meanRRs could create higher FDRs, but this affection was not very strong (Figure IB). These results were also similar for other situations: genes were associated with only first trait, only second trait, single traits (e.g., only first trait and both traits) (Figure SI, S2).

The correlation between simulated and estimated values of *π*_3_ was also assessed. For large meanRRs, high correlations were observed for all sample sizes. For smaller meanRRs (range here), *π*_3_ values were over-or underestimated (Figure S3). Flowever, these small differences were not much affected to main analyses (Figure 1, SI, S2).

#### 3.1.3 Power for single-trait risk gene discovery

We compared gene numbers identified by mTADA and extTADA using the same threshold PP > 0.8. For *π*_3_ = 0 (no overlapping information), mTADA and extTADA reported nearly the same positive gene numbers (Figure 2A). Flowever, mTADA identified more genes than extTADA when *π*_3_ increased. In addition, mTADA’s gene counts were also higher than those of extTADA when higher meanRRs were used.

**Figure 2:**
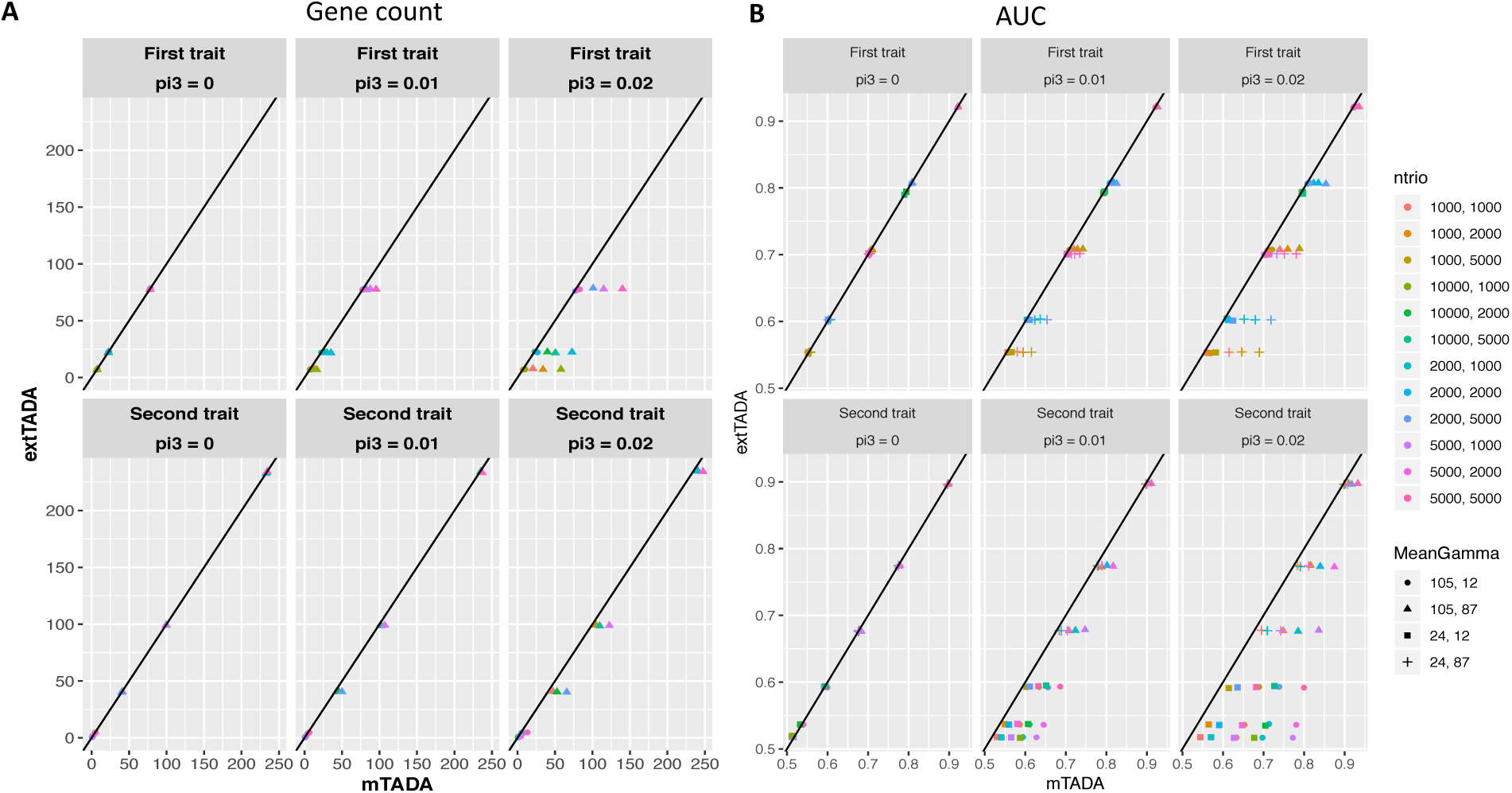
*Results of simulated data for single traits. Right panels show AUC* (*area under the Receiver Operating Characteristic* (*ROC*) *curve*) *results of mTADA and extTADA while left panels describe positive gene counts with PPs* > *0.8. MeanGamma*, *ntrio and pi3 describe for mean relative risks, sample sizes and the proportions of overlapping genes, ntrio and MeanGamma describe the information of two traits*.

#### 3.1.4 Comparison of AUCs for single traits

We designed a simulation experiment to assess the performance in the classification of risk and non-risk genes. We applied extTADA to single-trait data from our simulated data. We then calculated AUCs for mTADA and extTADA using classification results from single-trait data. AUCs of both were equal when *π*_3_ = 0 (Figure 2B). However, AUCs of mTADA were higher than those of extTADA when *π*_3_’s values were larger. In addition, mTADA also performed better extTADA with larger meanRRs.

### 3.2 Results of mTADA and extTADA on neuropsychiatric disease and CHD data

mTADA was applied to family data of 15 pairs of six disorders (ASD, SCZ, DD, ID, EPI and CHD). A threshold PP > 0.8 was used to prioritize top genes. To compare between mTADA and extTADA, we also extracted top prioritized genes from extTADA using the same threshold PP >

DD based results showed strong convergence with smaller credible intervals because of its large sample size as well as a high relative risks of DNMs (Figure 3). The highest *π* _3_ was observed for pairs of DD, ID, ASD and CHD (*π* _3_ > 0.019). These disorders also had the highest percentage of genes that overlap if we only focused on gene risk-gene proportions (Figure 3). CHD and EPI had the lowest *π*_3_, (0.001) followed by SCZ-EPI (0.0023). Figure S4 shows sampling results of the proportions of overlapping genes for each pair of these traits; and Table S 1 describes results of 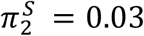, 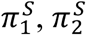 estimated by extTADA and *π* _3_ estimated by mTADA.

**Figure 3:**
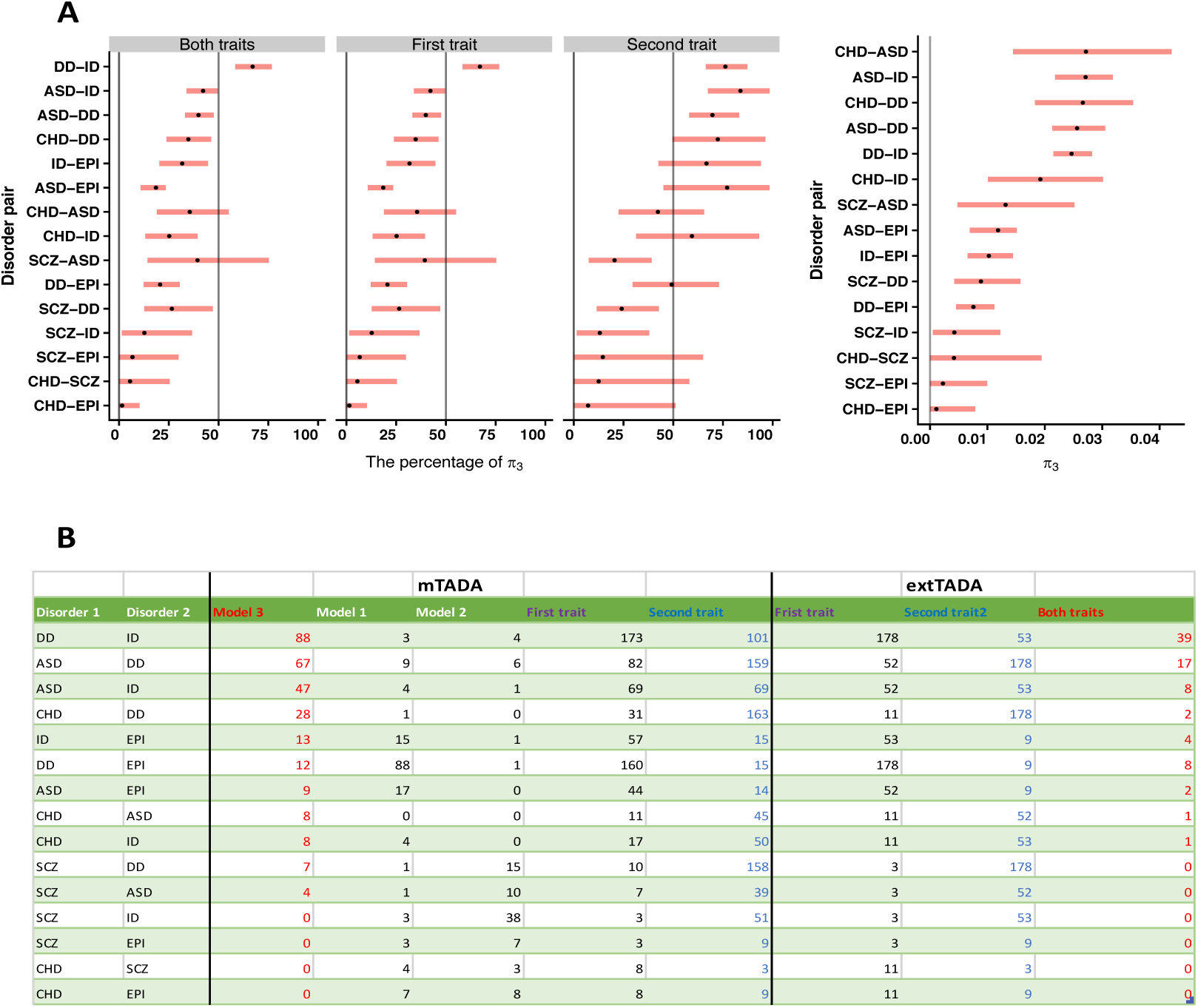
*Results of real data. A*) *Estimated results of the overlapping proportion of risk genes* (*tc*_3_) *for 15 pairs of 6 disorders: schizophrenia* (*SCZ*)*, congenital heart disease* (*CHD*)*, intellectual disability* (*ID*)*, developmental disorder* (*DD*)*, autism spectrum disorder* (*ASD*)*, epilepsy* (*EPI*)*. The first, second and third panels shows the proportion of risk genes that overlap for two traits* (*=π*_3_/(*π*_*1*_ + *π* _2_ + *π* _3_))*, first trait only* (*=π*_3_/(*π*_*1*_ *+ π*_3_)) *and second trait only* (*=π*_*3*_/(*π*_1_ + *π*_3_)) *while the fourth panel is π*_3_*. B*) *mTADA ‘s and extTADA ‘s results for all pairs of disorders by using posterior probabilities > 0.8. Numbers in cells are number of risk genes. For example*, *for the pair of DD and ID, mTADA’s results are: 3, 4 and 88 genes for Model 1, 2 and 3 respectively; 173 and 101 genes for DD and ID in that order*.

Regarding overlapping genes, the highest number was also observed for DD and ID (88 genes) followed by ASD-DD (67 genes) and ASD-ID (47 genes). Four pairs of traits (CHD-EPI, SCZ-EPI, CITD-SCZ, SCZ-ID) had no overlapping genes. In comparison with extTADA in the identification of overlapping genes, mTADA reported higher or equal gene numbers (Figure 3). Table S2 shows full mTADA’s results for these six disorders.

We also used mTADA to prioritize top genes for single traits and compared with extTADA. For DD and ID, mTADA always performed better extTADA (Figure 3). Similar results were also observed for ASD; except for the pair ASD-SCZ in which mTADA was better than extTADA for SCZ but extTADA was better than mTADA for ASD. For CHD, EPI and SCZ, mTADA was better than extTADA when CHD was combined with DD.

### Insights into top mTADA genes

#### Overlapping genes between two traits

Using a threshold PP>0.8, 153 genes were supported by the two-trait model in at least one pair (*π*_3_ > 0.8, Table S2). Seven genes (*ARID1B, GABRB3, KCNQ2, STXBP1, SYNGAP1, TLK2, POGZ, SCN2A*) were observed for at least six pairs of disorders (Table 2). POGZ and SCN2A were present in eight pairs of disorders POGZ was significant for pairs relating to ASD, DD, ID, CHD and SZ while SCN2A was significant for pairs relating to ASD, DD, EPI, ID and SCZ. We checked DNMs of these two genes. As expected, POGZ had no DNMs for CHD, and SCZN2A had no DNMs for EPI. Interestingly, in the latest CHD study (Jin, et al., 2017), POGZ was one of the top CHD gene while no DNMs were observed for SCN2A. In addition, in the latest study of 6,753 parent-offspring trios with NDDs and EPI (Heyne, et al., 2018), 16 DNMs were in POGZ, but only one DNM was from a patient who has both ID and EPI.

**Table 2:** *Genes with the posterior probabilities* (*PPs*) *of Model 3* (*two traits*) *> 0.8. These genes appear in at least 4 pairs of disorders. Cells shows the PP values.*

| Gene Name | ASD-DD | ASD-EPI | ASD-ID | CHD-ASD | CHD-DD | CHD-EPI | CHD-ID | CHD-SCZ | DD-EPI | DD-ID | ID-EPI | SCZ-ASD | SCZ-DD | SCZ-EPI | SCZ-ID |
| --- | --- | --- | --- | --- | --- | --- | --- | --- | --- | --- | --- | --- | --- | --- | --- |
| POGZ | 1.00 | 0.14 | 1.00 | 0.97 | 0.99 | 0.01 | 0.99 | 0.46 | 0.17 | 1.00 | 0.27 | 0.82 | 0.86 | 0.02 | 0.75 |
| SCN2A | 1.00 | 1.00 | 1.00 | 0.22 | 0.50 | 0.03 | 0.36 | 0.02 | 1.00 | 1.00 | 1.00 | 0.82 | 0.85 | 0.75 | 0.73 |
| ARID1B | 1.00 | 0.08 | 1.00 | 0.89 | 0.97 | 0.00 | 0.94 | 0.02 | 0.09 | 1.00 | 0.15 | 0.16 | 0.19 | 0.00 | 0.10 |
| GABRB3 | 0.98 | 0.99 | 0.94 | 0.28 | 0.61 | 0.06 | 0.28 | 0.00 | 1.00 | 0.98 | 0.99 | 0.14 | 0.20 | 0.14 | 0.06 |
| KCNQ2 | 0.82 | 0.87 | 0.90 | 0.05 | 0.66 | 0.05 | 0.51 | 0.00 | 1.00 | 1.00 | 1.00 | 0.02 | 0.22 | 0.13 | 0.11 |
| STXBP1 | 0.98 | 0.99 | 0.99 | 0.23 | 0.65 | 0.05 | 0.49 | 0.00 | 1.00 | 1.00 | 1.00 | 0.12 | 0.22 | 0.13 | 0.11 |
| TLK2 | 0.98 | 0.09 | 0.99 | 0.85 | 0.96 | 0.01 | 0.95 | 0.05 | 0.13 | 1.00 | 0.22 | 0.23 | 0.31 | 0.00 | 0.19 |
| CHD2 | 1.00 | 0.80 | 1.00 | 0.41 | 0.72 | 0.02 | 0.56 | 0.00 | 0.82 | 1.00 | 0.88 | 0.16 | 0.19 | 0.02 | 0.10 |
| CTNNB1 | 0.84 | 0.01 | 0.92 | 0.39 | 0.97 | 0.00 | 0.95 | 0.03 | 0.13 | 1.00 | 0.21 | 0.03 | 0.22 | 0.00 | 0.12 |
| MLL | 0.91 | 0.60 | 0.89 | 0.08 | 0.60 | 0.02 | 0.34 | 0.00 | 0.91 | 1.00 | 0.92 | 0.03 | 0.16 | 0.03 | 0.07 |
| SYNGAP1 | 1.00 | 0.08 | 1.00 | 0.32 | 0.63 | 0.00 | 0.48 | 0.03 | 0.09 | 1.00 | 0.15 | 0.83 | 0.86 | 0.01 | 0.74 |
| CACNA1A | 0.09 | 0.11 | 0.03 | 0.01 | 0.95 | 0.34 | 0.70 | 0.01 | 1.00 | 0.98 | 0.98 | 0.00 | 0.17 | 0.10 | 0.04 |
| KCNQ3 | 0.93 | 0.78 | 0.78 | 0.11 | 0.67 | 0.03 | 0.24 | 0.00 | 0.95 | 0.98 | 0.92 | 0.05 | 0.22 | 0.07 | 0.06 |
| MED13L | 0.90 | 0.01 | 0.95 | 0.19 | 0.82 | 0.00 | 0.72 | 0.01 | 0.06 | 1.00 | 0.09 | 0.03 | 0.20 | 0.00 | 0.10 |

#### Significant genes of single traits

To better understand mTADA results, we focused on three disorders (EPI, CHD and SCZ) whose DN-based genes have not been reported as many as the three other disorders. We used the *PP*_*ik*_ + *PP*_*i*3_ (*k = 1*, 2) of mTADA to obtain a single-trait’s PP for the *i*^*th*^ gene in the analysis of each pair of two disorders. We used DD as the main trait to infer results of other trait.

#### CHD

There were 31 genes with PP>0.8 by combining CHD and DD. 20/31 was not in the list of known CHD genes and in the meta-analysis results of a recent CHD study of Jin, et al. (2017) (Figure 4). We tested the PPIs of these 31 genes by using GeNets and STRING database. Based on GeNets, these genes were well connected to communities (Overall connectivity p < 2.2×10-e3, Figure 4). Next, by using STRING database, the number of edges were also higher than expected between these 31 nodes (20 edges versus 9 expected edges, p = 0.00198). We used expression data to test these genes. The majority of these genes were strongly expressed in early to late-mid prenatal stages (Figure 4).

**Figure 4:**
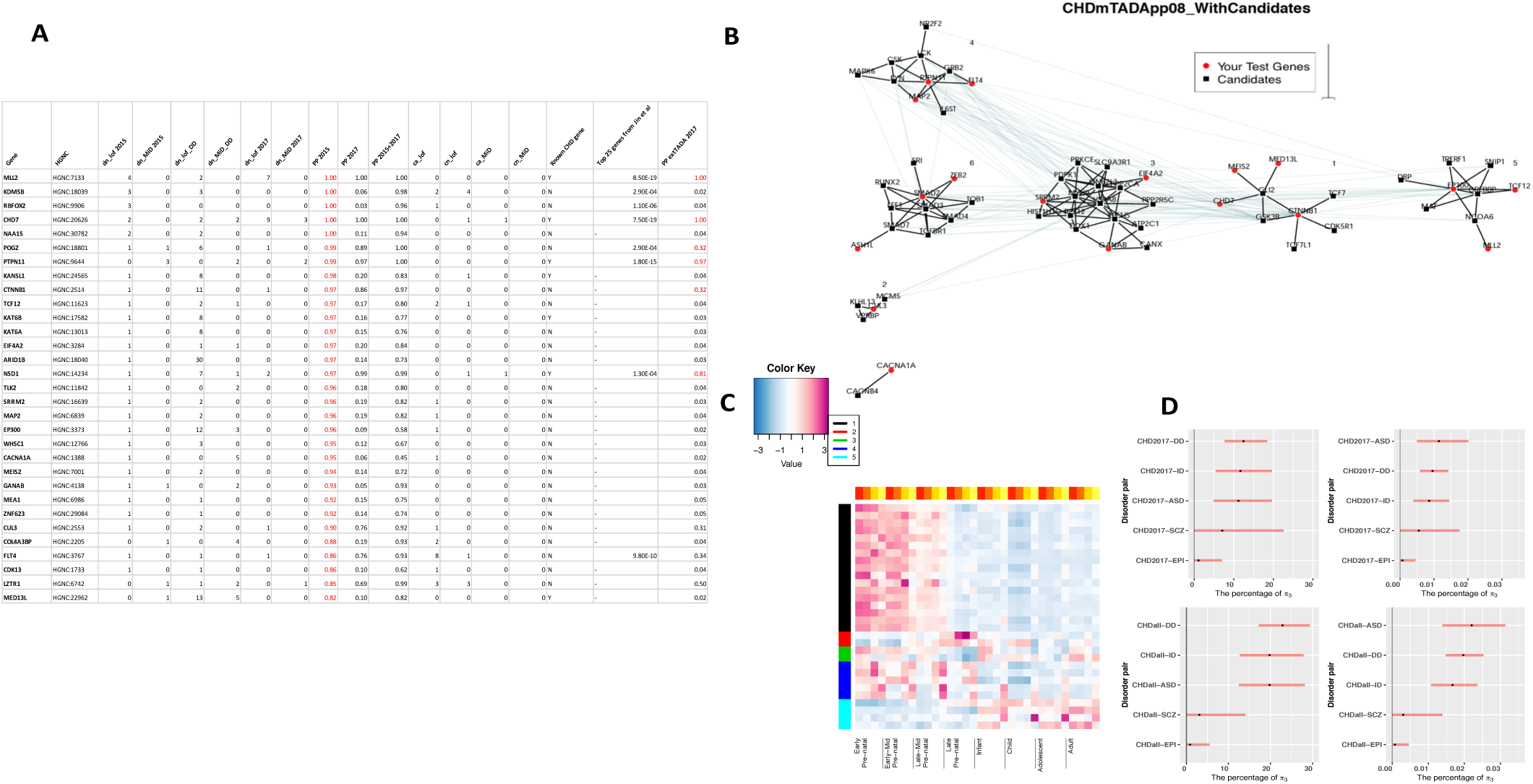
*Congenital heart disease* (*CHD*) *results by using developmental disorder* (*DD*) *information. This is the top 31 genes* (*posterior probabilities, PPs, > 0.8*) *identified by mTADA using the data set of Homsy, et al.* (*2015*). ***A***) *PPs of the 31 genes: dn J’of,MiD, ca JofiMiD, cn Jof/MiD describes loss-of-function/missense damaging de novo mutation, case and control variant counts respectively* (*the fourth and fifth columns are for DD data*). PP 2015, PP 2017, PP 2015+2017 describe PPs of CHD data from Homsy, et al. (*2015*), Jin, et al. (*2017*) *and both respectively; the sixteenth column describes whether a gene is a biown CHD gene* (*Y*) *or not* (*N*)*; the seventeenth column shows meta-p values calculated by Jin, et al.* (*2017*) *for their top 25 genes. The last column describes PPs of extTADA for an independent trio data of Jin, et al.* (*2017*). ***B*)** *The results of the analysis of protein-protein interactions.* ***C*)** *Results of the gene expression analysis.* ***D*)** *New mTADA results for the CHD data set of Jin, et al.* (*2017*): *top panels are for an independent data set while bottom panels are for full data sets.*

### Validation of CHD genes using an independent data set

We used the data of Jin, et al. (2017) to validate these results (See Methods). First, we tested the top CHD genes from the independent data set which include 1241 trios and 226 cases from Jin, et al. (2017). From the 1241 trios, three genes (CTNNB1, CUL3, LZTR1) of the 20 novel genes had LoF or MiD DNMs (p<2.00e-4, Figure 4). In addition, CUL3 had one LoF variant from case data. These three genes had only one DNMs in the primary analysis, and were not called as significant genes by extTADA. We also ran extTADA on the 1241 trios, and saw that 4 of the 31 genes had PP>0.8 (Figure 4). As expected, extTADA results of the majority of these 31 genes had low PPs because these genes had only 0 or 1 DNMs. Next, we compared our 31 genes with the top 25 genes meta-analyzed by Jin, et al. (2017). 8/31 were in the 25-gene list (p < 9.99e-05).

To better understand the performance of mTADA on the combination of CHD and other disorders, we applied mTADA to the independent data set only and the combined data set. First, we ran mTADA on the independent 1241-trio data. Similar to the primary analysis, we also saw high overlaps between CHD, ASD, DD and ID (Figure 4). By combining CITD and DD, 24 genes had PP>0.8. There were 6 overlapping genes between the 31 genes and the 24 genes (p < 9.99e-05). In these six genes, two genes NSD1 and CTNNB1 showed the benefit of using mTADA. Both genes had only one LoF DNM and did not show significant results in the original study of 1204 trios (Homsy, et al., 2015) as well as extTADA, but had highly significant PPs from mTADA (PP>0.97). Next, mTADA was applied to full 2451 trios. Estimated *π*_3_ values were similar to those of 1204 trios (Figure 4). There were 57 genes with PP>0.8.19/57 genes were in the 31 genes (p < 9.99e-05).

#### EPI

There were 15 genes with PP>0.8 by combining EPI and DD. Similar to top CHD genes, these genes were connected to communities (Overall connectivity p<2.2xl0-e3, Figure S5) by using GetNets. They also had more interactions than expected by using the STRING database (22 edges versus 2 expected edges, p = l.lle-05). Four genes GABBR2, HECW2, MLL, WDR19 were not in the list of known EPI genes. GABBR2 was reported as a top gene by extTADA (PP=0.97), but HECW2, MLL, WDR19 had only PP<0.3 in extTADA. Interestingly, both GABBR2, HECW2 had DNMs in a WGS study recently (Hamdan, et al., 2017). We also used expression data to test these EPI genes. Differently from CITD, these genes were expressed in different stages of the human brain (Figure S5).

#### SCZ

There were 10 genes with PP>0.8 including AUTS2, BRPF1, HIST1H1E, MAP4K4, MKI67, POGZ, SCN2A, SETD1A, SYNGAP1, TAF13. These were significantly connected to communities by the PPI analysis (Overall connectivity p<2.2xl0-e3, Figure S6) by using GeNets. These genes were not strongly connected by using STRING database (1 edges versus 0 expected edges, p = 0.22). In these genes, only TAF13 and SETD1A were suggested as top genes in previous study (Fromer, et al., 2012; Nguyen, et al., 2017). In addition, AUTS2 (PP>0.8) was reported as a SCZ genes from a common variant based study (Zhang, et al., 2014). Using expression data, multiple genes of these genes were expressed in prenatal stages, but the signals were as not strong as those of CHD (Figure S6).

## 4. Discussion

In this paper, we propose a method to jointly analyze two traits (mTADA) using *de novo* exome sequencing data. The method is an extension of our previous work for single traits (He, et al., 2013; Nguyen, et al., 2017). mTADA estimates the proportion of overlapping genes between two traits, and then uses this information to infer how many overlapping genes exist between two traits. The pipeline is also able to infer the number of risk genes for each trait by calculating posterior probabilities (PPs) of genes for each trait. On simulated data, mTADA performs better than extTADA on the identification of risk genes. We also applied mTADA to more than 13,000 trios of five disorders and reported overlapping genes between these disorders. We also saw that mTADA reported more risk genes for these disorders than extTADA. This suggests that mTADA can help in the identification of additional risk genes, especially for disorders whose large sample sizes are challenging to obtain or whose mean relative risks are small. For such disorders, users can combine the data of the disorders with large public data sets (e.g., trio data of ASD, DD) to prioritize risk genes. Using one-trait information to leverage the information for other traits has been successful in fine-mapping (Kichaev, et al., 2017) and common-variant (Maier, et al., 2018) studies. Based on our best knowledge, mTADA is the first tool using this approach for de novo mutation data We hope that mTADA (https://github.com/hoangtn/mTADA) will be generally useful for analyzing *de novo* mutation data across complex traits.

By using mTADA, multiple overlapping genes were observed for CHD, DD, ID and ASD. This replicates a recent study (Homsy, et al., 2015) in which high overlapping genes were observed for CFID and neurodevelopmental disorders (NDDs). Interestingly, CHD did not show any overlapping information with another NDD: epilepsy (EPI). Two genes SCN2A and POGZ which have been reported as risk genes for some of these disorders (Stessman, et al., 2016; White, et al., 2016; Ben-Shalom, et al., 2017; Jin, et al., 2017) are top overlapping genes from mTADA (Table S2), but they show different trends. No SCN2A DNMs are in CFID data and no POGZ DNMs are in EPI data. One possible reason is that the sample size of EPI is small in this study (356 trios). Another hypothesis might be because they do not have strong overlapping biological pathways. We did not see any overlapping information between SCZ and CHD as well as SCZ and CHD. We analyzed in depth the top prioritized genes of CHD, EPI and SCZ (Figure 4, S5-6). Top risk CFID and EPI genes from mTADA are also reported in recent study (Hamdan, et al., 2017; Jin, et al., 2017). Multiple top CFID genes have only one DNM, but have DNMs or variants in independent data sets (Figure 4). This suggests that they might be real risk-genes for this disorder. Interesting, we identify 20 novel CHD genes (posterior probabilities > 0.8) and 3 of these 20 genes have DNMs in an independent data set. This shows the benefit of using mTADA in the prediction of risk genes for CHD by borrowing the information of DD. CHD genes are strongly expressed in pre-natal stages of the human brain (Figure 4).

Although mTADA performs better than the single-trait based extTADA, it does have some limitations. mTADA uses the parameters of single traits from extTADA to infer the proportion of overlapping genes (*π*_3_). Using parameters from extTADA makes mTADA much faster in its calculation, it means mTADA relies on the results of the single-trait pipeline extTADA that uses a full Bayesian approach. Also, mTADA as well as extTADA use de novo counts for each gene and divide these counts into different categories similar to other rare variant based studies (Sanders, et al., 2012; De Rubeis, et al., 2014; Genovese, et al., 2016; Allen, et al., 2017). Future studies which are able to incorporate the annotation information of each DNM may increase the power of mTADA or similar tools.

With further development, the mTADA approach can be generalized further to consider more than two traits simultaneously, and the increased information could increase the number of identified risk genes but at a cost of increased computational time.

In conclusion, mTADA can be very useful for better understanding the genetic correlation across disorders (via the proportion of overlapping genes), and to prioritize additional risk genes for disorders. The approach of mTADA can be used to identify shared/specific risk genes for different categories of one trait (e.g., loss of function and missense *de novo* mutations). Genetic information of de novo mutations and rare case/control variants can be different (Sifrim, et al., 2016), mTADA might be adopted to pipelines which are able to apply to DNMs and rCCVs as two traits.

## Author’s contributions

Designed the pipeline used in analysis; performed the experiments, analyzed the data and drafted the manuscript: HTN. Conceived and designed the experiments: HTN, EAS. Contributed reagents/materials/analysis tools: HTN, AD, DP, JB, XH, PFS, EAS. Wrote the paper: HTN, AD, PFS, XH, EAS.

## Acknowledgements

This work is supported by NTH grant R01MH105554 to E.A.S, and by Nil I grant R01MH110555 to D.P. The Sweden exome sequencing data generation and analysis are supported by the Stanley Center for Psychiatric Research and NIH grant R01 MH077139 to P.F.S. This work was supported in part through the computational resources and staff expertise provided by Scientific Computing at the Icahn School of Medicine at Mount Sinai. We are deeply grateful for the participation of all subjects contributing to this research.

## References

Allen, A.S., et al. Ultra-rare genetic variation in common epilepsies: a case-control sequencing study. The Lancet Neurology 2017;16(2):135–143.

Allison, D.B., et al. Multiple phenotype modeling in gene-mapping studies of quantitative traits: power advantages. Am J Hum Genet 1998;63(4):1190–1201.

Barber, M.J., et al. Genome-wide association of lipid-lowering response to statins in combined study populations. PLoS One 2010;5(3):e9763.

Ben-Shalom, R., et al. Opposing Effects on NaV1.2 Function Underlie Differences Between SCN2A Variants Observed in Individuals With Autism Spectrum Disorder or Infantile Seizures. Biol Psychiatry 2017;82(3):224–232.

Carpenter, B., et al. Stan: A probabilistic programming language. Journal of Statistical Software 2016;20(2):1–37.

De Rubeis, S., et al. Synaptic, transcriptional and chromatin genes disrupted in autism. Nature 2014;515(7526):209–215.

Deciphering Developmental Disorders Study. Prevalence and architecture of de novo mutations in developmental disorders. Nature 2017;542(7642):433–438.

Fromer, M., et al. Discovery and statistical genotyping of copy-number variation from whole-exome sequencing depth. Am J Hum Genet 2012;91(4):597–607.

Fromer, M., et al. De novo mutations in schizophrenia implicate synaptic networks. Nature 2014;506(7487):179–184.

Galesloot, T.E., et al. A comparison of multivariate genome-wide association methods. PLoS One 2014;9(4):e95923.

Genovese, G., et al. Increased burden of ultra-rare protein-altering variants among 4,877 individuals with schizophrenia. Nature neuroscience 2016;19(11):1433.

Giambartolomei, C., et al. Bayesian test for colocalisation between pairs of genetic association studies using summary statistics. PLoS genetics 2014;10(5):e1004383.

Hamdan, F.F., et al. High Rate of Recurrent De Novo Mutations in Developmental and Epileptic Encephalopathies. Am J Hum Genet 2017;101(5):664–685.

He, X., et al. Integrated model of de novo and inherited genetic variants yields greater power to identify risk genes. PLoS Genet 2013;9(8):e1003671.

Heyne, H.O., et al. De novo variants in neurodevelopmental disorders with epilepsy. Nat Genet 2018;50(7):1048–1053.

Hoischen, A., Krumm, N. and Eichler, E.E. Prioritization of neurodevelopmental disease genes by discovery of new mutations. Nature neuroscience 2014;17(6):764.

Homsy, J., et al. De novo mutations in congenital heart disease with neurodevelopmental and other congenital anomalies. Science 2015;350(6265):1262–1266.

Iossifov, I., et al. The contribution of de novo coding mutations to autism spectrum disorder. Nature 2014;515(7526):216–221.

Jin, S.C., et al. Contribution of rare inherited and de novo variants in 2,871 congenital heart disease probands. Nat Genet 2017;49(11):1593–1601.

Kichaev, G., et al. Improved methods for multi-trait fine mapping of pleiotropic risk loci. Bioinformatics 2017;33(2):248–255.

Kucukelbir, A., et al. Automatic variational inference in Stan. In, Advances in neural information processing systems. 2015. p. 568–576.

Lelieveld, S.H., et al. Meta-analysis of 2,104 trios provides support for 10 new genes for intellectual disability. Nat Neurosci 2016;19(9):1194–1196.

Li, J., et al. Genes with de novo mutations are shared by four neuropsychiatric disorders discovered from NPdenovo database. Molecular psychiatry 2016;21(2):290.

Li, T., et al. A scored human protein-protein interaction network to catalyze genomic interpretation. Nat Methods 2017;14(1):61–64.

Lin, G.N., et al. Spatiotemporal 16p11.2 protein network implicates cortical late mid-fetal brain development and KCTD13-Cul3-RhoA pathway in psychiatric diseases. Neuron 2015;85(4):742–754.

Loader, C. Locfit: Local regression, likelihood and density estimation. R package version 2007;1.

Lutz, S.M., et al. A general approach to testing for pleiotropy with rare and common variants. Genetic epidemiology 2017;41(2):163–170.

Maier, R.M., et al. Improving genetic prediction by leveraging genetic correlations among human diseases and traits. Nat Commun 2018;9(1):989.

Miller, J.A., et al. Transcriptional landscape of the prenatal human brain. Nature 2014;508(7495):199–206.

Nguyen, H.T., et al. Integrated Bayesian analysis of rare exonic variants to identify risk genes for schizophrenia and neurodevelopmental disorders. Genome Med 2017;9(1):114.

Pickrell, J.K., et al. Detection and interpretation of shared genetic influences on 42 human traits. Nature genetics 2016;48(7):709–717.

Sanders, S.J., et al. De novo mutations revealed by whole-exome sequencing are strongly associated with autism. Nature 2012;485(7397):237.

Sifrim, A., et al. Distinct genetic architectures for syndromic and nonsyndromic congenital heart defects identified by exome sequencing. Nat Genet 2016;48(9):1060–1065.

Solovieff, N., et al. Pleiotropy in complex traits: challenges and strategies. Nat Rev Genet 2013;14(7):483–495.

Stessman, H.A.F., et al. Disruption of POGZ Is Associated with Intellectual Disability and Autism Spectrum Disorders. Am J Hum Genet 2016;98(3):541–552.

Szklarczyk, D., et al. The STRING database in 2017: quality-controlled protein-protein association networks, made broadly accessible. Nucleic Acids Res 2017;45(D1):D362–D368.

Takata, A., et al. De Novo Synonymous Mutations in Regulatory Elements Contribute to the Genetic Etiology of Autism and Schizophrenia. Neuron 2016;89(5):940–947.

Turley, P., et al. Multi-trait analysis of genome-wide association summary statistics using MTAG. Nature genetics 2018:1.

White, J., et al. POGZ truncating alleles cause syndromic intellectual disability. Genome Med 2016;8(1):3.

Willsey, A.J., et al. The Psychiatric Cell Map Initiative: A Convergent Systems Biological Approach to Illuminating Key Molecular Pathways in Neuropsychiatric Disorders. Cell 2018;174(3):505–520.

Zhang, B., et al. Association study identifying a new susceptibility gene (AUTS2) for schizophrenia. Int J Mol Sci 2014;15(11):19406–19416.

Zhernakova, A., et al. Meta-analysis of genome-wide association studies in celiac disease and rheumatoid arthritis identifies fourteen non-HLA shared loci. PLoS Genet 2011;7(2):e1002004.

